# Drought is a stronger driver of plant morphology and nutritional composition than warming in two common pasture species: Effects of drought and warming on pasture quality

**DOI:** 10.1101/2021.02.16.431509

**Authors:** Karen L. M. Catunda, Amber C. Churchill, Haiyang Zhang, Sally A. Power, Ben D. Moore

## Abstract

Under warmer and drier future conditions, global livestock and dairy production are threatened by impacts on the productivity and nutritional quality of pastures. However, morphological and nutritional adjustments within plants in response to warming and drought vary among species and less is known how these relate to production and forage quality. To investigate this, we grew two common pasture species, tall fescue (*Festuca arundinacea*: grass) and lucerne (*Medicago sativa*: legume), in a climate-controlled facility, under different temperatures (ambient and elevated) and watering regimes (well-watered and droughted). We found that drought had a strong negative impact on biomass production, morphology and nutritional quality while warming only significantly affected both species when response metrics were considered in concert, although to a lesser degree than the drought. Furthermore, interactions between warming and drought were only seen for lucerne, with the greatest reduction in biomass and most dead material and dry matter content. In tall fescue, drought had bigger impacts on nutritional composition than morphological traits, while in lucerne, drought affected all morphological traits and most nutritional parameters. These findings suggest that in future climate scenarios, drought may be a stronger driver of changes in the morphology and nutritional composition of pasture grasses and legumes, compared to modest levels of warming.

## 1 INTRODUCTION

As the climate changes, more extreme and frequent periods of heat stress and water deficiency have become the most common and critical limiting factors for productivity and nutritional quality of pastures and grasslands across the globe (Chang-Fung-Martel et al., 2017; Deléglise et al., 2015; IPCC, 2014). Grazing livestock require a reliable and consistent supply of high-quality forage (Herrero et al., 2013; Lee et al., 2017) to achieve high animal performance and maintain profitable production (Dairy Australia, 2018; Laca et al 2001). However, research suggests that pasture systems will be challenged to meet demands for forage in some regions of the world as predicted climate change will impact annual pasture production by driving shifts in plant phenology and increases the inter-annual variability of production. Both of these outcomes pose an increased climate risk to the dairy and meat industries (Perera et al., 2020; Rojas-Downing et al., 2017). Along with changes in pasture productivity, there are also potential shifts in the morphological traits and nutritional composition of pasture species that are likely to alter forage quality and digestibility (AbdElgawad et al., 2014; Herrero et al., 2015; Howden et al., 2008). In contrast to shifts in productivity, these changes and their consequences under future climate scenarios, such as warming and drought, are relatively understudied.

Warming can affect plant growth directly, with the nature of the response dependent upon the optimal temperature of a plant species. For example, positive warming effects on growth can be expected for species where production is limited by cold temperatures (Bloor et al., 2010), whereas neutral or negative responses may be more likely in warmer environments (Dukes et al., 2005). Warming above a plant’s optimal temperature may cause temperature stress via direct effects on plant physiology and metabolism and indirectly via increased evapotranspiration and lowered plant water availability (Rustad et al., 2001). These stresses can negatively influence pasture forage quality via affect morphological traits, such as reduced leaf size, tiller emergence and leaf:stem ratios (Mitchell, 1956; Wilson et al., 1991). In addition, some studies have reported a reduction in the nutritional quality of forage under warming (Lee et al., 2017) through increases in concentrations of fibre, and decreases in concentrations of crude protein (Waghorn & Clark, 2004) and non-structural carbohydrates (Habermann et al., 2019; Wilson et al., 1991).

Drought and its associated reduction in soil moisture content result in a wide range of impacts on plant morphology and nutritional composition, with the magnitude of impacts dependent on the developmental stage of plants and the severity and duration of the drought (Gray & Brady, 2016; IPCC, 2014). Severe drought inhibits growth and accelerates maturation of existing plant tissue, death of tillers and leaf senescence that result in decreasing the leaf:stem ratio and increasing in fibre concentrations (Bruinenberg et al., 2002; Ren et al., 2016). Additionally, severe water deficit increases nutrient translocation (mainly nitrogen and non-structural carbohydrates) from leaves to roots, thus reducing the concentrations of nutrients aboveground (Buxton, 1996; Durand et al., 2009). In contrast, moderate drought stress typically induces different morphological responses including delays in plant maturation and growth especially for perennial species, which then results in only mild to moderate leaf loss (Buxton, 1996). Studies focused on nutritional responses have reported seemingly idiosyncratic changes among different nutritional parameters such as no effect or reductions in fibre concentrations, and no effect or slight improvements in crude protein concentrations and digestibility of forage under moderate water deficit conditions (Bittman, 1988; Buxton, 1996; Deleglise et al., 2015; Dumont et al., 2015; Kuchenmeister et al., 2013; Staniak & Harasim, 2018). In these cases, unchanged or reduced fibre concentrations could be explained by reduced growth and increases in leaf:stem ratio reported under moderate conditions (Bruinenberg et al., 2002; Deleglise et al., 2015). Increased crude protein concentrations could be attributed to delayed maturation and lower biomass production under moderate water limitation, allowing nutrients such as nitrogen to become concentrated in plant tissues (Dumont et al., 2015; Grant et al., 2014).

While studies have been conducted on responses of species to a single climate change variable, there is a lack of information as to what may happen with concurrent changes in temperature and water availability (Chang-Fung-Martel et al., 2017; Viciedo et al., 2021). In addition, how morphological and nutritional adjustments can affect each other in different plant species under climate change scenarios, especially how warming and drought interact within pasture species, are not well understood and need to be investigated. To address these research gaps, we conducted a study in a climate-controlled facility to investigate the effects of warming and short-term drought, alone and in combination, on plant morphological traits and nutritional composition of two important temperate pasture species: tall fescue (*Festuca arundinaceae* – a C_3_ grass) and lucerne (*Medicago sativa* – a C_3_ legume). These species were chosen as they are perennial forage species, widely used in many countries throughout the world. We grew plants under different average daily temperatures [ambient: 26 °C; elevated: 30 °C] and watering regimes [well-watered: 100% soil water holding capacity (WHC); droughted: 40% WHC]. We hypothesized that warming and drought, as isolated factors, will negatively affect biomass production, morphological traits and nutritional quality, with the magnitude dependent on species-specific responses. We further predict that drought in combination with warming will have a more pronounced negative impact on plant morphological and nutritional traits than as isolated climate factors, as warming can exacerbate the negative effects of water stress in the plants.

## 2 MATERIALS AND METHODS

### 2.1 Site and experimental design

The experiment was conducted between mid-April and early-August 2018 in four climate-controlled glasshouse chambers located at the Hawkesbury Campus of Western Sydney University, in Richmond, NSW, Australia (33°36’40’’ S, 150°44’43’’ E). Two agriculturally important temperate pasture species were used in this experiment: the C_3_ grass, tall fescue (*Festuca arundinacea*) and the C_3_ legume, lucerne (*Medicago sativa)*. The two-factor experimental design included two temperatures (ambient, aT; elevated, eT) and two watering regimes (well-watered, W; droughted, D), giving four treatment combinations (aT.W, aT.D, eT.W, eT.D), each with eight replicate pots and thus 32 pots per species. Treatment replicates were divided evenly among four chambers, with two chamber replicates maintained at aT and the other two at eT. Watering regimes were nested within the temperature treatments (Zhang et al., 2021). Pot positions within chambers were re-randomized every two weeks to minimize potential within-chamber effects.

### 2.2 Plant growth conditions and treatments

We collected field soil (sandy-loam texture with a soil pH ∼5.6) from the Pastures and Climate Extremes (PACE) field experimental facility, also located on the Western Sydney University Hawkesbury Campus (33°36’S,150°44’E) (Churchill et al., 2020). The soil was sieved (5 mm), air-dried and mixed with quartz sand (7:3, v/v) then 3.9 kg of soil was placed into each plastic pot (3.7 L, 150 mm diameter, 240 mm height). Seeds of each species were surface-sterilized with a 1.25% NaOCl for 10 min, rinsed with deionized water 10 times and germinated in Petri dishes with sterilized water for 1 week. Five germinated seedlings were transplanted into each pot and then thinned to four healthy individuals per pot after 2 weeks (Zhang et al., 2021). The legume pots were supplied with appropriate rhizobia (Easy Rhiz soluble legume inoculant, Group AL, New Edge Microbials, NSW, Australia) necessary for nodulation (Zhang et al., 2021).

The temperature treatments included an ambient regime (aT; target of 26/18 °C day/night) and an elevated temperature regime (warming) with + 4 °C warming (eT; target of 30/22 °C day/night) using a 15:9 light: dark cycle. The ambient regime reflected the average daily maximum temperature for the site over the previous 20 years (Australian Government Bureau of Meteorology, Richmond RAAF site), and the elevated regime represented the predicted increase of 4 °C for this region within this century (Pearce et al., 2007). Temperature treatments were initiated at the same time as transplantation. Humidity was controlled at 60% and the effectiveness for achieving the temperature conditions within the glasshouse chambers are reported in Zhang et al. (2021).

To maintain consistent water availability, an automated irrigation system was used to ensure pots were well-watered every second day to 100% water holding capacity (WHC, i.e. until drainage just occurred after watering events). All pots were maintained at well-watered conditions until the two watering regimes (well-watered and droughted) were initiated three weeks before the final harvest. Pots in well-watered treatments were maintained in well-watered condition as before. In contrast, pots in drought treatments had watering withheld for 4 days (until the majority of the plants started to wilt). Thereafter, we weighed pots every other day to maintain the drought treatments pots at 40% WHC for one week (adding an appropriate amount of water when necessary). After one week, we re-watered the drought treatments pots to bring the soil water condition back to 100% WHC. This oscillating drought regime (shifting between 100% and 40% WHC) was repeated for the remaining three weeks of the experiment, ending just prior to the final harvest. Within each watering regime, pots under aT and eT were maintained at similar WHC, to be able to test the direct effects of warming and minimize the interactive influence of warming on soil water content. Following typical pasture management practices, we also implemented an aboveground biomass clipping event followed by nutrient fertilization in our experiment. Plant shoots were clipped 8 weeks after planting (before imposing the drought treatment) at 5 cm above the soil surface, and allowed to regrow until the final harvest. Two weeks after the clipping, we applied fertilizer (containing KNO_3_ and KH_2_PO_4_ resulting in a fertilizer pulse of 30 kg N ha^-1^ and 5 kg P ha^-1^; Zhang et al., 2021).

### 2.3 Harvest and morphological traits measurements

Immediately prior to harvest, plant height and number of tillers/stems per pot were measured for both species. Plant height (cm) was measured in each pot from ground level to the tip of the tallest plant; subsequently, the numbers of tillers (grass) and stems (legume) from each pot were counted. The percentage of standing aboveground biomass that was dead was estimated visually and assigned to five classes (0%, 25%, 50%, 75%, 100%) ranging from the entire plant being healthy to all aboveground plant tissue being senesced. To harvest aboveground biomass, plants were cut at the level of the soil surface, biomass was weighed, immediately frozen (−18°C) and later freeze-dried, then weighed to determine the total dry biomass (g/ pot). Dry biomass of lucerne was sorted into leaves (plus flowers when present) and stems, and fractions were weighed to calculate the leaf:stem ratio. Dry leaves of tall fescue could not be distinguished from tillers.

### 2.4 Sample processing

Dried biomass samples were ground to pass through a 1-mm screen in a laboratory mill (Foss Cyclotec Mill, Denmark) and stored in airtight plastic containers in the dark at room temperature prior to collection of near-infrared reflectance spectra and wet chemistry analysis. Prior to nitrogen analysis, subsamples of biomass from each pot were reground to a fine powder using a ball-mill (Retsch® MM200; Hann, Germany). Nutritional analysis was performed on the total aboveground harvested material (a mixture of leaves, stems/tillers and flowers when present), including a mixture of both live and dead material.

### 2.5 Nutritional analysis

All dried and ground samples were each scanned twice and their spectra were collected and averaged using near-infrared reflectance spectroscopy (NIRS; FOSS XDS Rapid Content™ Analyzer) with a spectral range from 400 to 2500 nm. Half of the samples for each species (4 replicates per treatment and per species) were selected for determining nutrient composition by wet chemistry for all parameters, except for dry matter and crude protein contents (for which all samples were analysed). Samples for wet chemistry were selected using the ‘select’ function in the software WinISI 4.8.0 (FOSS Analytical A/S, Denmark) to represent the range of spectral variation in the population, summarized by a principal component analysis to minimize redundancy in spectra (Catunda et al., unpublished manuscript).

For wet chemistry, the selected samples were subjected to analyses of dry matter (DM) and ash (ASH) according to the standard methods and procedures for animal feed outlined by the Association of Official Analytical Chemists (AOAC, 1990). Nitrogen (N) concentration was determined from ∼ 100 mg samples using an automated combustion method on a Leco TruMac CN-analyzer (Leco Corporation, USA). Crude protein (CP) concentration was then calculated by applying a 6.25 conversion factor to the N concentration (AOAC, 1990). Ether extract (EE) was determined according to the American Oil Chemists’ Society-AOCS high-temperature method using petroleum ether and the Soxhlet method (Buchi 810 Soxhlet Multihead Extract Rack, UK). Fibre fractions were determined with an ANKOM Fibre Analyzer (model 200, ANKOM® Technology, NY, USA) with use of neutral and acid detergent solutions and corrected for dry matter content (Goering & Van Soest, 1970). Samples were analysed for neutral detergent fibre (NDF), acid detergent fibre (ADF) and acid detergent lignin (ADL) by the sequential method of Van Soest & Robertson (1980). Sodium sulphite and α-amylase were added to the solution for NDF determination. The concentration of hemicellulose (HEM) was calculated by the difference between NDF and ADF concentrations after sequential analysis, while cellulose (CEL) concentration was calculated as the difference between ADF and ADL. The values of ASH, EE, CP and NDF were used to calculate non-structural carbohydrates (NSC) according to Sniffen et al. (1992). Estimated digestible dry matter (DDM), expressed as percentage of dry matter, was calculated according to *Equation 1* below (Linn & Martin, 1989):

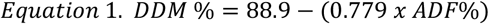

For the development of NIRS calibration models, modified Partial Least Squares regression with cross-validation was used to develop predictive equations for each nutritional parameter to prevent overfitting of models (Shenk & Westerhaus 1991). Standard normal variate and detrend mathematical transformations (Barnes et al., 1989) were applied to raw NIR spectra to reduce the influence of particle size, and a variety of derivative mathematical pre-treatments were employed to decrease spectral noise effects. The best predictive models were selected on the basis of the highest coefficient of determination of calibration (R^2^) and the internal cross-validation (one minus the variance ratio, 1-VR), and the lowest standard error of calibration (SEC) and internal cross-validation (SECV), and the smallest difference between SEC and SECV (Catunda et al., unpublished manuscript; Norman et al., 2020). The best models developed for each nutritional parameters were used to predict the other half of the samples. The mathematical treatment of spectra and descriptive statistics for NIRS calibrations can be found in **Table A1**.

### 2.7 Calculations and statistical analysis

Plant biomass, morphological traits and nutritional composition (expressed as a percentage of dry matter) were analysed statistically using linear mixed-effects (LME) models in the ‘lme4’ package in R (Bates et al., 2015). Temperatures (T), watering regimes (W), and their interactions (T x W), were defined as fixed effects and the glasshouse chambers were specified as a random factor. Residuals were checked for normality and we applied log-transformation to the percentage of dead material as a continuous response. Data for each species were analysed separately and we were not explicitly interested in contrasting the two species. We calculated the mean effect size due to drought (response ratio) based on the ratio of each mean value in the drought treatments (D) to the mean value in the well-watered (W) treatments at each temperature level (aT *= Equation 2*; eT *= Equation 3*) along with 95% confidence intervals (CI). In the effect size figures, positive values represent responses that are greater under drought than in well-watered treatments for the respective temperature level, while negative values represent the opposite.

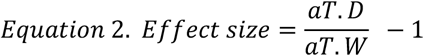

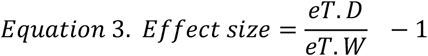

To achieve a more holistic overview of changes brought about by climate treatments on plant response variables for each pasture species that accounts for the non-independence of within-plant chemistry, we performed a multivariate, principal components analysis (PCA). To test for the effects of both temperatures and watering regimes on plant biomass, morphological and nutritional responses, we undertook permutational analysis of variance (PERMANOVA) using the ‘vegan’ package in R. All statistical analyses were carried out using the software R version 3.5.2 (R Core Team, 2019).

## 3 RESULTS

### 3.1 Impacts of warming and short-term drought on plant biomass and morphological responses

Drought, but not warming, significantly affected plant dry biomass or morphological traits for both species (**Table A2, Table 1**). For tall fescue, drought significantly decreased biomass by 24% (*p* < 0.01, **Figure 1A**) and increased percentage of dead material by 19% (*p* < 0.01, **Figure 2A**). There was, however, no effect on plant height or number of tillers (**Figure 1A**). For lucerne, drought significantly (*p* < 0.01 for all parameters) decreased biomass (51%, **Figure 1B**), plant height (18%), number of stems (28%) and leaf:stem ratio (40%), as well as increased the percentage of dead material (21%, **Figure 2B**). Overall, the negative effect of drought on biomass and morphological traits were stronger in lucerne than that in tall fescue.

**Table 1.**
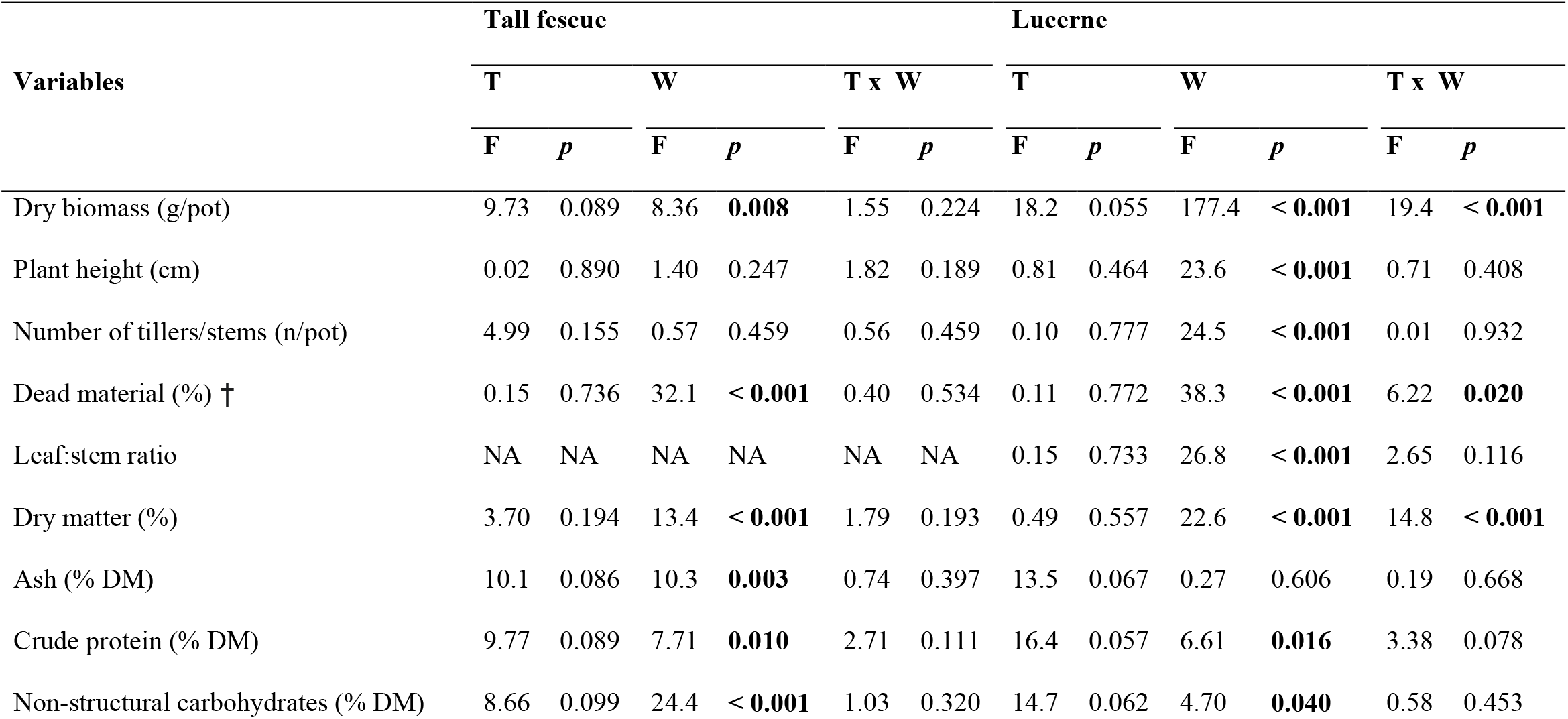
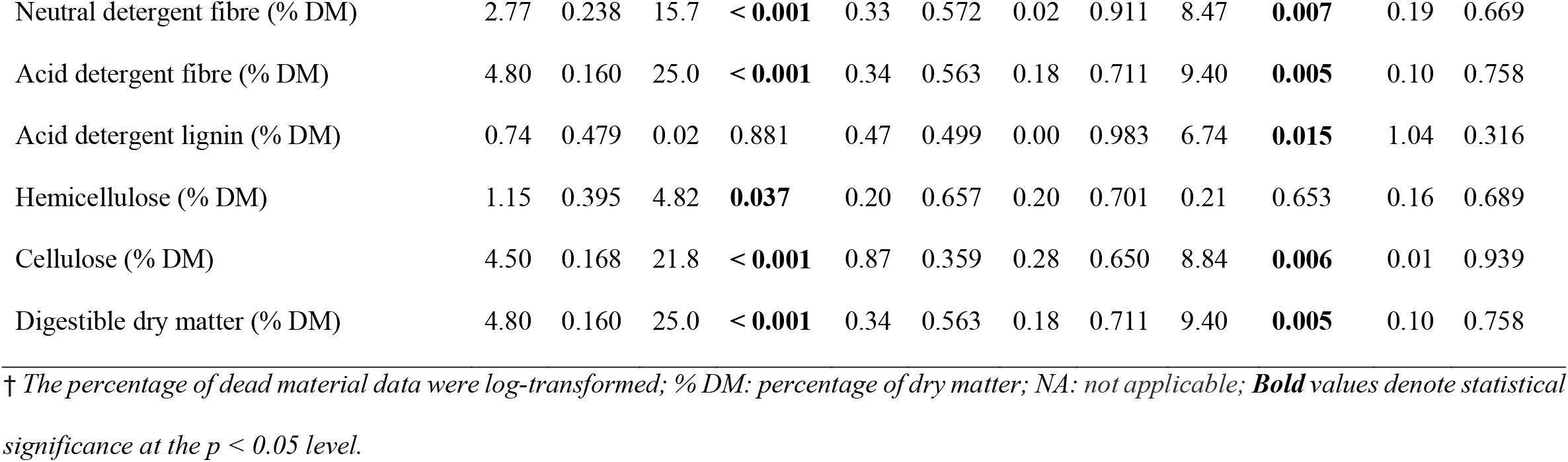
Linear mixed effects models with fixed effects for temperatures (T), watering regimes (W) and their interaction (T x W) for plant dry biomass, morphological traits and nutritional parameters of tall fescue (*Festuca arundinacea*) and lucerne (*Medicago sativa*). Growth chamber was included as a random effect.

**Figure 1.**
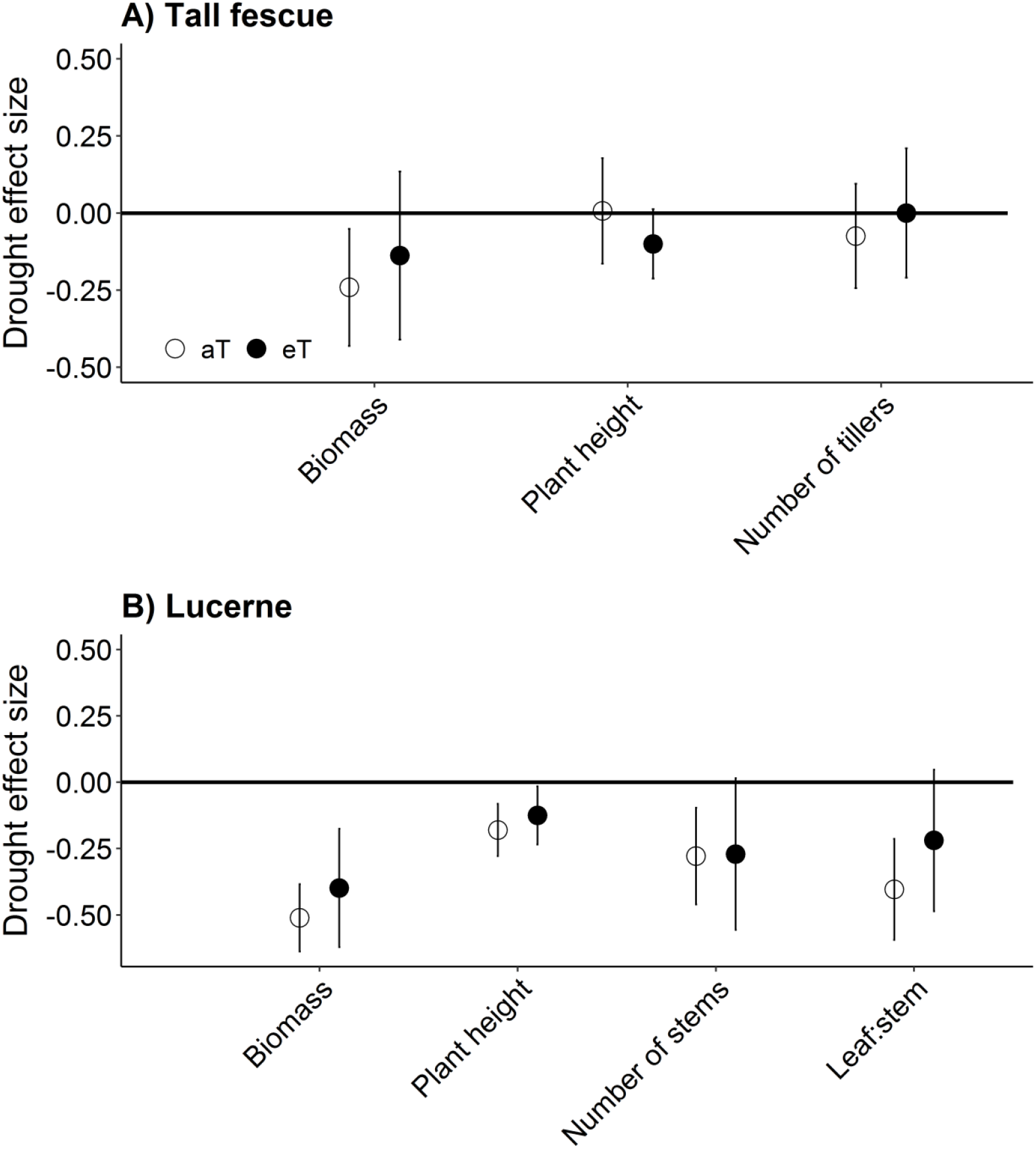
Drought effect sizes under ambient (aT, open circle) and elevated (eT, closed circle) temperatures on plant dry biomass and morphological traits of A) tall fescue (*Festuca arundinacea*) and B) lucerne (*Medicago sativa*). Values shown are means with vertical bars representing 95% confidence intervals (n = 8).

**Figure 2.**
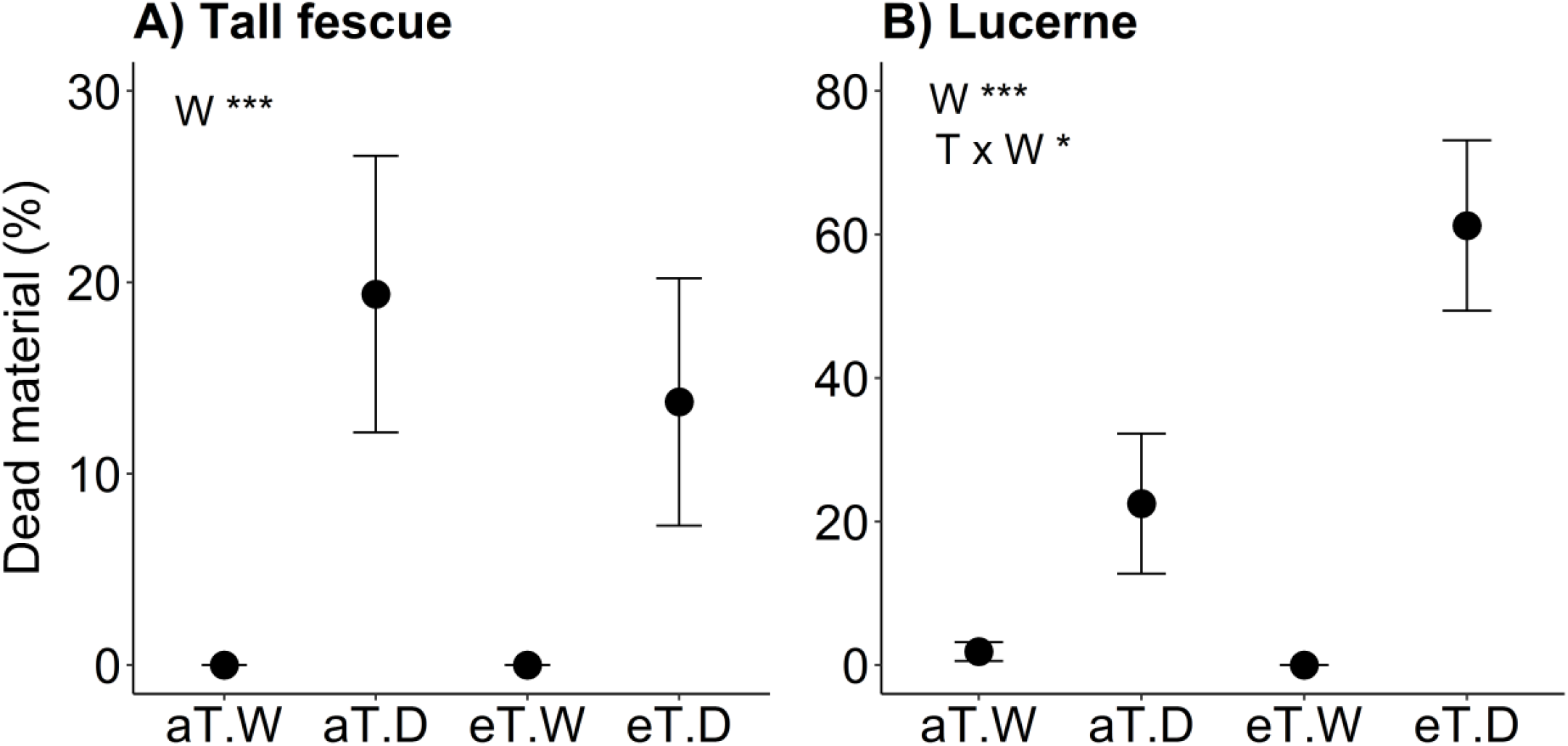
Percentage of dead material (%) for A) tall fescue (*Festuca arundinacea*) and B) lucerne (*Medicago sativa*) grown under different temperatures (ambient, aT; elevated, eT) and watering regimes (well-watered, W; droughted, D). Values shown are mean ± 1 standard error (n = 8). Treatment codes for indicating significance for fixed effects are as follows: T= temperatures, W= watering regimes, and T x W their interaction. Significance levels as follows: * *p* < 0.05, *** *p* < 0.001.

There was not a significant interaction between temperatures and watering regimes (T x W) for biomass or morphological traits in tall fescue, although, for lucerne, the interaction significantly affected biomass (*p* < 0.01, **Figure 1B**) and the percentage of dead material (*p* = 0.02, **Figure 2B**). Specifically, for lucerne, warming (eT) partially offset the negative effect of drought on biomass (−40%, **Figure 1B**), but exacerbated its effect on the percentage of dead material (+60%, **Figure 2B**).

### 3.2 Impacts of warming and short-term drought on nutritional responses

Drought, but not warming, significantly affected nutritional parameters for both species (**Table A2, Table 1**). For tall fescue, drought significantly affected all parameters of nutritional composition, except acid detergent lignin (ADL, *p* = 0.88). Drought resulted in a significant decrease in non-structural carbohydrates (NSC, 12% **Figure 3A**, *p* < 0.01) and digestible dry matter (DDM, 3%, *p* < 0.01), and a significant increase in dry matter (DM, 7%, *p* **<** 0.01), ash (ASH, 15%, *p* < 0.01), crude protein (CP, 9%, *p* = 0.01), neutral detergent fibre (NDF, 7% *p* < 0.01), acid detergent fibre (ADF, 11%, *p* < 0.01), hemicellulose (HEM, 4%, *p* = 0.04) and cellulose (CEL, 11%, *p* < 0.01).

**Figure 3.**
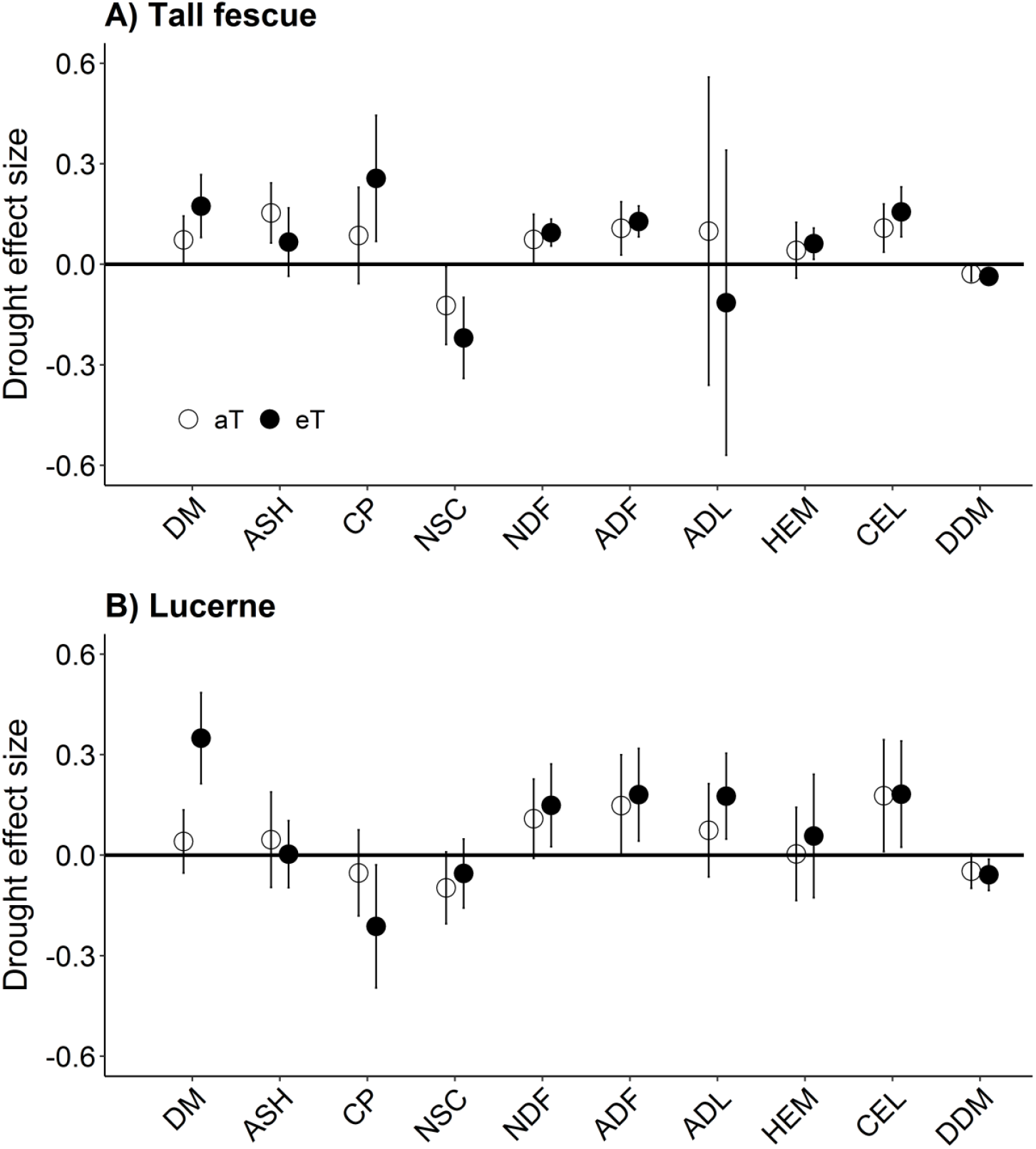
Drought effect sizes under ambient (aT, open circle) and elevated (eT, closed circle) temperatures on nutritional parameters of A) tall fescue (*Festuca arundinacea*) and B) lucerne (*Medicago sativa*). Abbreviations are as follows: dry matter (DM), ash (ASH), crude protein (CP), non-structural carbohydrates (NSC), neutral detergent fibre (NDF), acid detergent fibre (ADF), acid detergent lignin (ADL), hemicellulose (HEM), cellulose (CEL) and digestible dry matter (DDM). Values shown are means with vertical bars representing 95% confidence intervals (n = 8).

For lucerne, drought significantly affected all nutritional parameters, except ASH (*p* = 0.61) and HEM (*p* = 0.65). In contrast to its effects in tall fescue, drought significantly decreased CP by 5% in lucerne (**Figure 3B**, *p* = 0.02) and caused a significant 7% increase in ADL (*p* = 0.02). In addition, compared to the aT.W treatment, drought significantly decreased NSC and DDM by 10% (*p* = 0.04) and 5% (*p* < 0.01), respectively, while significantly increasing DM, NDF, ADF, and CEL by 4% (*p* < 0.01), 11% (*p* < 0.01), 15% (*p* < 0.01) and 18% (*p* < 0.01), respectively.

There was no significant interaction between temperatures and watering regimes for nutritional parameters in either species (*p* > 0.05). However, the combination of warming and drought increased the DM of lucerne by 35% (*p* < 0.01), compared with ambient temperature and well-watered plants.

### 3.3 Assessing plant biomass, morphological and nutritional responses to warming and drought in a multivariate context

We used a multivariate approach to assess overall plant responses to the climate treatments. For tall fescue we found that both temperature (PERMANOVA: *p* = 0.018, **Figure 4**) and watering regime (PERMANOVA: *p* = 0.001) significantly influenced plant responses, but there was no interaction (T x W) between treatments (PERMANOVA: *p* = 0.600). However, for lucerne we found that temperature (PERMANOVA: *p* = 0.008, **Figure 5**), watering regime (PERMANOVA: *p* = 0.001) and their interaction (PERMANOVA: *p* = 0.020) significantly influenced plant responses. The ellipses in **Figure 4A** and **Figure 5A** show statistically significant treatment separation in the trait-space for both species. In tall fescue (**Figure 4A**), the most significant separation of these plant responses occurred when under both warming and drought scenarios, while in lucerne (**Figure 5A**), the driver separating plant responses were mainly from drought rather than warming.

**Figure 4.**
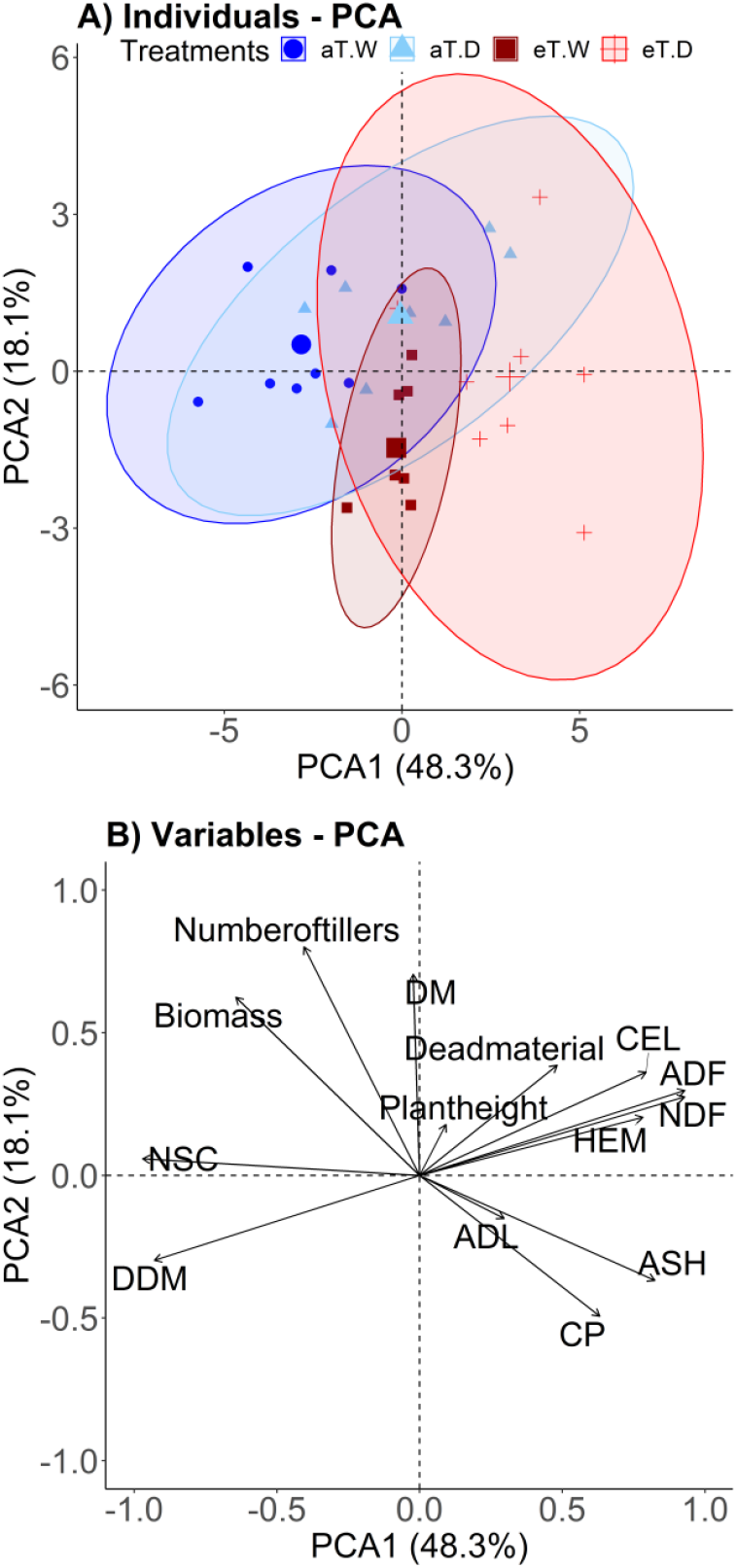
Principal component analysis biplot illustrating A) scores for plant individuals, grouped by treatment (ambient, aT; elevated, eT, well-watered, W; droughted, D) with 95% confidence ellipses (PERMANOVA: Temperatures *p* = 0.018, Watering regimes *p* = 0.001, Temperatures x Watering regimes *p* = 0.600) and B) variables loadings for tall fescue (*Festuca arundinacea*). Morphological traits include plant height, number of tillers and percentage of dead material. The parameters of nutritional composition follow the abbreviations in Figure 3. The symbol shape and colour of each point correspond to climate treatments.

**Figure 5.**
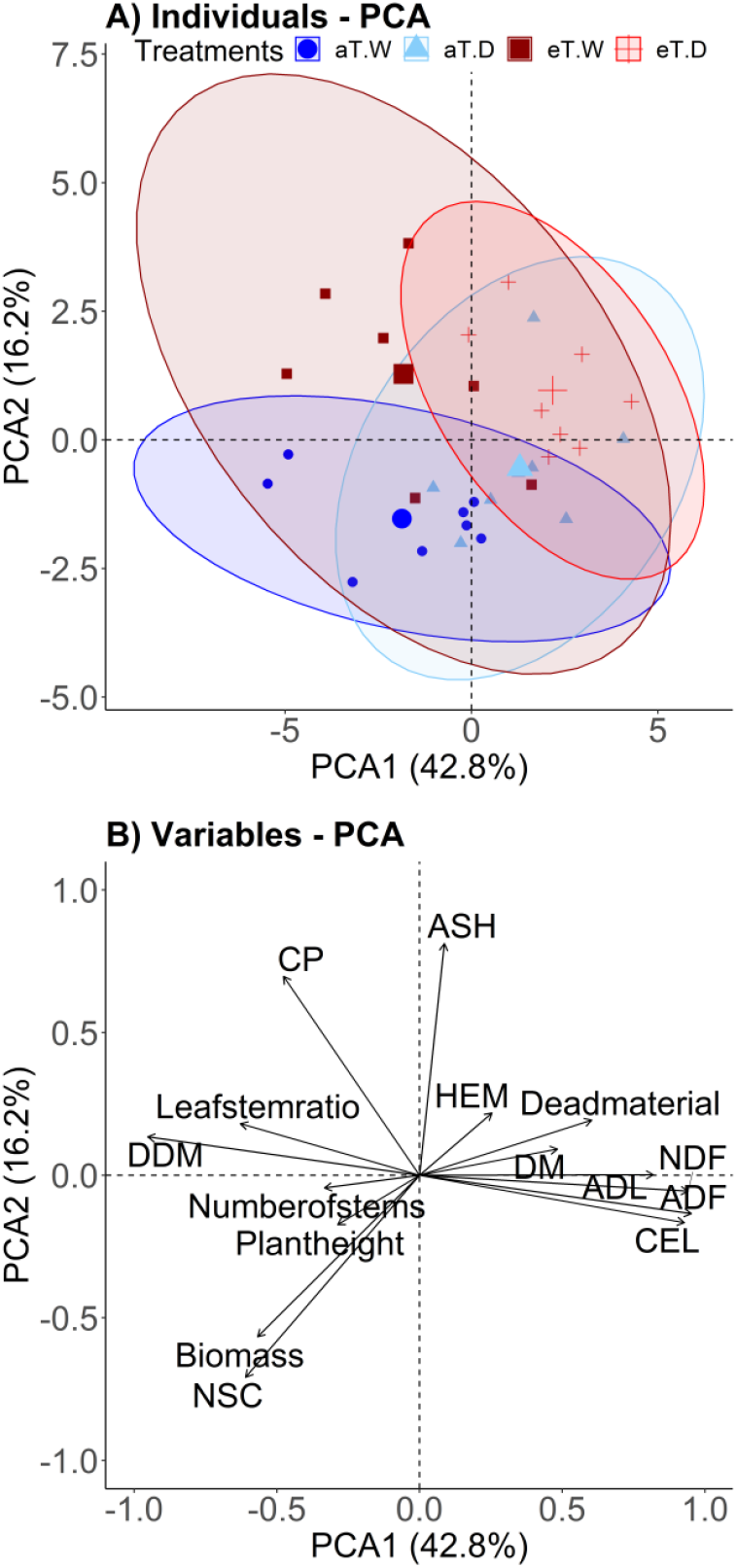
Principal component analysis biplot illustrating A) scores for plant individuals, grouped by treatment (ambient, aT; elevated, eT, well-watered, W; droughted, D) with 95% confidence ellipses (PERMANOVA: Temperatures *p* = 0.008, Watering regimes *p* = 0.001, Temperatures x Watering regimes *p* = 0.020) and B) variables loadings for lucerne (*Medicago sativa*). Morphological traits include plant height, number of stems, percentage of dead material and leaf:stem ratio. The parameters of nutritional composition follow the abbreviations in Figure 3. The symbol shape and colour of each point correspond to climate treatments.

In tall fescue, the first two principal components explained 66% of the variation across treatments (**Figure 4B**). The first principal component (PCA1, explaining 48.3% of the data variance) had positive loadings of fibre (NDF, ADF) and ASH, and negative loadings with DDM and NSC (**Figure 4B**). The second PCA axis (PCA2, explaining 18.1% of the data variance) was associated with plant morphological traits and nutritional parameters, with positive loadings for biomass and number of tillers, and negative loadings for CP. In addition, we found that biomass and the number of tillers were negatively correlated with CP, while the percentage of dead material was positively correlated with fibre fractions (ADF particularly) and consequently negatively correlated with DDM (**Figure 4B, Figure A1**).

In lucerne, PCA1 and PCA2 explained respectively 42.8% and 16.2% of variation in sample biomass, morphological traits and nutritional parameters (**Figure 5B**). PCA1 (**Figure 5B**), had positive loadings for fibre (NDF, ADF, CEL, ADL) and percentage of dead material, and negative loadings for DDM and leaf:stem ratio. PCA2 was associated with higher CP and ASH, and negatively associated with biomass and NSC. Lucerne biomass is positively correlated with NSC and negatively correlated with fibre (CEL particularly), and that fibre (CEL particularly) is negatively correlated with leaf:stem ratio and positively correlated with the percentage of dead material (**Figure 5B, Figure A2**).

In addition, for both pasture species (**Figure 4, Figure 5**) the biomass, DDM and NSC were higher for those plants grown under aT.W treatment, while high fibre concentrations and a greater percentage of dead material were associated with the eT.D treatment. Particularly in lucerne (**Figure 5**), eT.W treatment was associated with high leaf:stem ratio and CP. Overall, PCA1 showed that the nutritional parameters explained the highest percentage of variance in tall fescue (**Figure 4B**) while both the morphological traits and nutritional parameters explained the variance in lucerne (**Figure 5B**).

## 4 DISCUSSION

Here, we have determined the effects of warming, short-term drought and, their interaction on plant biomass, morphological traits and nutritional composition of two common pasture species (tall fescue and lucerne). We found that exposure of these species to short-term drought resulted in a significant negative impact on biomass production, morphology and nutritional quality; warming did not significantly affect individual growth or nutritional parameters but did have a significant overall effect on both species when assessed in a multivariate context. In addition, we found a significant interaction between warming and drought in lucerne, which resulted in greater differences between well-watered and drought treatments at elevated than at ambient temperature. These findings demonstrate that drought had far bigger impacts than warming overall, the effects of warming were greater when combined with drought – conditions that more closely reflect predicted climates under which grazing systems will function in the future.

### 4.1 Impacts of warming on plant biomass, morphological and nutritional responses

For both pasture species we found limited evidence of shifts in plant biomass, morphological, or nutritional responses associated with continuous warming. Previous studies have shown species-specific positive (Bloor et al., 2010; Dieleman et al., 2012), negative (Cantarel et al., 2013; Lee et al., 2017) and neutral (Dukes et al., 2005; Dumont et al., 2015) warming effects on productivity and/or quality in forage species associated with regional climatic differences. A widely anticipated mechanism by which warming can indirectly affect plants is via increased evapotranspiration and consequent reductions in soil water content (Rustad et al., 2001). In our experiment the eT.W and aT.W treatments were maintained at similar WHC, to be able to isolate the direct effects of air temperature, while minimizing the interactive influence of warming on soil water content. This approach and resultant lack of significant warming effects on biomass, morphological traits and nutritional composition suggests that evapotranspiration-mediated indirect effects might be responsible for many of the observed changes in forage productivity and quality attributed to warming under field conditions (Cantarel et al., 2013). In general, in our study warming was also the minor contributor to interaction effects with drought. Our results suggest that if soil water availability can be maintained under field conditions, then it may be possible to minimise the anticipated negative impacts of rising temperatures on forage nutritional quality, at least for the species and temperatures in this study, if not more widely.

### 4.2 Impacts of short-term drought on plant biomass, morphological and nutritional responses

In our experiment, short-term drought significantly decreased biomass while increasing the percentage of dead material for both species, and particularly for lucerne, negatively affecting all morphological traits. In tall fescue, the short-term drought did not alter plant height and the number of tillers, instead, drought influenced plant phenology through accelerated senescence of existing plant tissue. In particular for lucerne, the negative effect of drought on plant height and leaf:stem ratio can be explained by water deficiency having reduced plant growth and accelerated senescence, resulting in relatively more stem material compared to leaves, as also reported in previous studies with pasture species (Bruinenberg et al., 2002; Ren et al., 2016). These findings suggest that morphological changes in lucerne play a major role in plant adaptation responses under drought stress. Overall, the morphological responses found in our study, suggest that these responses must be related to different abilities to tolerate and adapt to the drought that is species-specific (Lee et al., 2013).

In our study, although drought produced different morphological changes in these two species, a decrease in nutritional quality and digestibility was common to both. The significant reductions in NSC and increases in fibre (especially cellulose) may be explained by the high percentage of dead material and for lucerne decreases in leaf:stem ratio that were found in this study. In support of our results, previous studies found that stems are associated with more fibre, higher forage toughness and consequently lower digestibility (Bruinenberg et al., 2002; Buxton, 1996; Durand et al., 2009; Ren et al., 2016). Here, some nutritional responses to drought did differ between species, for example, in terms of fibre fractions such as hemicellulose (increased only in tall fescue) and lignin (increased only in lucerne) concentrations. According to previous studies with forage species, the differences in fibre fractions responses to drought between species may reflect differences in plant structure (Amiri et al., 2012; Pontes et al., 2007). For example, a study showed that legume stem tissue is thick with a high bulk density and is comprised of a considerably larger undegradable fraction like lignin compared to grasses species (Amiri et al., 2012). This may have contributed to the bigger drought response on the lignin concentration of lucerne found in our study. Supporting this, lignin concentrations found in our study in lucerne were four times those found in tall fescue.

Were also found species differences in crude protein responses to drought. In tall fescue, CP concentration increase under drought can be explained by trade-offs between concentration and growth, such that lower biomass production increased the tissue concentration of plant CP, as also reported by previous studies investigating the water stress effects on forage quality in grasslands (Dumont et al., 2015; Grant et al., 2014). In lucerne, the decrease found in CP under drought can be linked to reduced nitrogen fixation and/ or lower nutrient uptake. Studies on forage legumes have found that drought-stressed plants reduce the biological nitrogen fixation activity for root-associated rhizobia (Kuchenmeister et al., 2013; Liu et al., 2018; Zahran, 1999). Additionally, a meta-analysis of forage species observed that plant nutrient uptake was lower under dry soils (Dumont et al., 2015). Under severe drought, reduced nutrient uptake is typically driven by reduced diffusion of nutrients through the soil as well as reduced root ability to transfer nutrients to aboveground tissue, thereby contributing to lower CP concentrations (Evans & Burke, 2013; Durand et al., 2009; Gonzalez-Dugo et al., 2005). Finally, previous studies have reported that senescence of aboveground materials promotes nutrient translocation (mainly nitrogen and soluble carbohydrates) from leaves to roots (Durand et al., 2009; Buxton, 1996), which may explain in our study the reduction of CP in lucerne, and NSC in both species, under drought conditions.

### 4.3 Assessing plant biomass, morphological and nutritional responses to warming and drought in a multivariate context

By adopting a multivariate framework to capture a holistic view of plant responses, we detected significant effects of both warming and drought and, for lucerne, a significant interaction effect. The latter was seen as a strengthening of warming effects under drought. This additive effect of warming on drought treatment in lucerne is in line with results reported by other recent studies who also showed the additive effect on plant growth and nutrients responses (Dellar et al., 2018; Orians et al., 2019). Multivariate analysis is not widely applied in agricultural research, particularly in feed evaluation, however, our findings suggest that this oversight may underestimate the consequences of climate change for forage quality. Statistical ordination techniques like PCA can usefully reduce the complexity of large forage data sets, aiding interpretation (Gallo et al., 2013; Pezzopane et al., 2020) while also avoiding the issue of multiple comparisons posed by numerous univariate analyses and non-independence of the chemical constituents in individual plants. In this study, for tall fescue, the majority of variability under climate change treatments was first associated (PCA1) with nutritional parameters and secondarily (PCA2) associated with morphological parameters. For lucerne, shifts in morphological and nutritional parameters contributed similarly to treatment differences. Our findings suggested that nutritional composition should be an essential component of studies aimed at evaluating the impacts of climate change on pasture species. Although in this study the responses of individual compounds were informative, our multivariate analyses of pasture morphology and nutritional quality provided a more comprehensive perspective of climate change impacts for future field conditions where the effects of multiple factors occur simultaneously across many aspects of plant biology and nutritional chemistry.

In conclusion, drought, even in the short term, can be a strong driver of change in many individual morphological traits and the nutritional composition of pasture species. However, when considered in a multivariate framework, warming also has a significant impact on plant morphology and nutritional quality, although to a lesser degree than the drought. We found that exposure of these pasture species to warmer and drier conditions resulted in significantly less forage produced and a decline in nutritional quality. Furthermore, the potential negative impacts on nutritional quality will have implications for pasture species choice, animal production, and methane emissions. Improved understanding of changes in morphology which might, in turn, affect forage quality among several pasture species under climate change can lead to more efficient use of resources, better economic outcomes, consequently, an improvement in the future of sustainable livestock production around the globe.

## ACKNOWLEDGEMENTS

We are grateful to all the team members in Pasture and Climate Extreme (PACE) project specially Chioma Igwenagu, Gil Won Kim, Kathryn Fuller, Lena Schmidt, and Vinod Jacob. This work was supported by funding from the Meat & Livestock Australia Donor Company and Dairy Australia. The authors would like to thank the technical team at Western Sydney University for technical support. The authors declare that there are no conflicts of interest regarding the publication of this article.

