## APPENDIX for "Drought is a stronger driver of plant morphology and nutritional composition than warming in two common pasture species: Effects of drought and warming on pasture quality"

**Table A1**. Details of calibration sets, treatment of spectra, and regression statistics for NIRS modified partial least-squares regression models predicting nutritional composition parameters for tall fescue (*Festuca arundinacea*) and lucerne (*Medicago sativa*).

| Species | Parameter | Math treatment* | Calibration | | | | Validation | |
|---|---|---|---|---|---|---|---|---|
| | | | SEC | $R^2$c | SECV | 1-VR | SEP | $R^2$v |
| Tall fescue | ASH | 3,10,10,1 | 0.35 | 0.98 | 0.87 | 0.87 | 0.73 | 0.93 |
| | EE | 2,4,4,1 | 0.43 | 0.95 | 0.80 | 0.83 | 0.98 | 0.75 |
| | NDF | 2,8,6,1 | 1.78 | 0.98 | 2.50 | 0.96 | 2.34 | 0.97 |
| | ADF | 2,4,4,1 | 0.83 | 0.98 | 1.67 | 0.92 | 1.38 | 0.95 |
| | ADL | 3,10,10,1 | 1.14 | 0.85 | 1.51 | 0.73 | 1.55 | 0.71 |
| Lucerne | ASH | 2,8,6,1 | 0.41 | 0.96 | 0.73 | 0.88 | 0.91 | 0.89 |
| | EE | 2,6,4,1 | 0.54 | 0.92 | 0.89 | 0.79 | 0.99 | 0.72 |
| | NDF | 2,8,6,1 | 1.78 | 0.98 | 2.50 | 0.96 | 2.34 | 0.97 |
| | ADF | 2,4,4,1 | 1.41 | 0.94 | 1.80 | 0.89 | 2.06 | 0.91 |
| | ADL | 3,10,10,1 | 1.14 | 0.85 | 1.51 | 0.73 | 1.55 | 0.71 |

*The scatter correction used was standard normal variate + detrend and the range wavelengths used was 400-2492 nm for all parameters. Parameters expressed in % of dry matter: ASH: ash, EE: ether extract, NDF: neutral detergent fibre, ADF: acid detergent fibre, and ADL: acid detergent lignin. *Math treatment describes the mathematical treatment applied to the spectra (stored as log(1/reflectance)). The first two numbers describe the derivative used, the third and fourth numbers indicate the degrees of primary and secondary smoothing performed on the derivative. Thus 2,4,4,1 indicates that the second derivative was calculated with a gap size of 4 nm and that a maximal primary smooth but no secondary smooth was used. SEC: standard error of calibration; $R^2$c: coefficient of determination of calibration; SECV: standard error of* *the internal cross validation; 1-VR: coefficient of determination of the internal cross-validation; SEP: standard error of prediction; $R^2$v: coefficient of determination of validation.*

**Table A2.** Mean values (n = 8) and standard errors (SE) for plant dry biomass, morphological traits and nutritional parameters of tall fescue (*Festuca arundinacea*) and lucerne (*Medicago sativa*) grown under different temperatures (ambient, aT; elevated, eT) and watering regimes (well-watered, W; droughted, D).

| **Variables** | **Tall fescue** | | | | | | | | **Lucerne** | | | | | | | |
|---|---|---|---|---|---|---|---|---|---|---|---|---|---|---|---|---|
| | **aT.W** | | **aT.D** | | **eT.W** | | **eT.D** | | **aT.W** | | **aT.D** | | **eT.W** | | **eT.D** | |
| | **Mean** | **SE** | **Mean** | **SE** | **Mean** | **SE** | **Mean** | **SE** | **Mean** | **SE** | **Mean** | **SE** | **Mean** | **SE** | **Mean** | **SE** |
| Dry biomass (g/pot) | 9.64 | 0.55 | 7.31 | 0.57 | 6.69 | 0.43 | 5.76 | 0.71 | 25.9 | 0.63 | 12.6 | 0.76 | 16.9 | 0.58 | 10.1 | 1.10 |
| Plant height (cm) | 55.3 | 3.28 | 55.7 | 3.58 | 59.6 | 2.95 | 53.6 | 1.55 | 72.6 | 2.53 | 59.5 | 2.16 | 75.0 | 1.97 | 65.6 | 3.24 |
| Number of tillers/stems (n/pot) | 30.1 | 2.01 | 27.9 | 1.53 | 21.4 | 1.43 | 21.4 | 1.79 | 25.1 | 1.98 | 18.1 | 0.89 | 27.4 | 2.56 | 20.0 | 2.24 |
| Dead material (%) | 0 | 0 | 19.4 | 7.22 | 0 | 0 | 13.7 | 6.46 | 1.87 | 1.31 | 22.5 | 9.77 | 0 | 0 | 61.2 | 11.9 |
| Leaf:stem ratio | NA | NA | NA | NA | NA | NA | NA | NA | 1.09 | 0.04 | 0.65 | 0.06 | 1.01 | 0.06 | 0.79 | 0.10 |
| DM (%) | 31.7 | 0.69 | 34.0 | 1.00 | 28.5 | 0.55 | 33.4 | 1.46 | 32.6 | 0.73 | 34.0 | 1.44 | 30.8 | 0.94 | 41.5 | 2.58 |
| ASH (% DM) | 6.31 | 0.19 | 7.29 | 0.24 | 8.39 | 0.19 | 8.95 | 0.42 | 6.16 | 0.32 | 6.44 | 0.33 | 7.40 | 0.25 | 7.42 | 0.29 |
| CP (% DM) | 3.48 | 0.17 | 3.78 | 0.20 | 4.57 | 0.29 | 5.74 | 0.42 | 13.4 | 0.74 | 12.7 | 0.44 | 18.6 | 1.43 | 14.7 | 0.78 |

| | | | | | | | | | | | | | | |
|---|---|---|---|---|---|---|---|---|---|---|---|---|---|---|
| NSC (% DM) | 41.7 | 1.42 | 36.6 | 1.78 | 35.4 | 0.66 | 27.7 | 1.62 | 42.3 | 1.49 | 38.1 | 1.60 | 35.7 | 1.69 | 33.7 | 0.76 |
| NDF (% DM) | 48.4 | 1.33 | 52.0 | 1.40 | 50.6 | 0.61 | 55.4 | 0.92 | 38.2 | 2.02 | 42.4 | 1.23 | 37.7 | 2.19 | 43.3 | 1.06 |
| ADF (% DM) | 23.8 | 0.67 | 26.4 | 0.76 | 25.3 | 0.42 | 28.5 | 0.46 | 27.8 | 1.88 | 32.0 | 1.20 | 28.0 | 1.83 | 33.0 | 0.87 |
| ADL (% DM) | 1.97 | 0.20 | 2.17 | 0.46 | 2.66 | 0.29 | 2.36 | 0.48 | 7.93 | 0.49 | 8.52 | 0.29 | 7.54 | 0.48 | 8.86 | 0.14 |
| HEM (% DM) | 24.6 | 0.78 | 25.6 | 0.73 | 25.3 | 0.32 | 26.9 | 0.54 | 10.4 | 0.49 | 10.4 | 0.56 | 9.70 | 0.82 | 10.3 | 0.42 |
| CEL (% DM) | 21.9 | 0.59 | 24.2 | 0.60 | 22.6 | 0.53 | 26.2 | 0.78 | 19.9 | 1.45 | 23.4 | 1.03 | 20.5 | 1.50 | 24.2 | 0.81 |
| DDM (% DM) | 70.3 | 0.52 | 68.3 | 0.59 | 69.2 | 0.33 | 66.7 | 0.36 | 67.2 | 1.46 | 64.0 | 0.93 | 67.1 | 1.43 | 63.2 | 0.68 |

*% DM: percentage of dry matter; DM: dry matter; CP: crude protein; NSC: non-structural carbohydrates; NDF: neutral detergent fibre; ADF: acid detergent fibre; ADL: acid detergent lignin; HEM: hemicellulose; CEL: cellulose; DDM: digestible dry matter; NA: not applicable.*

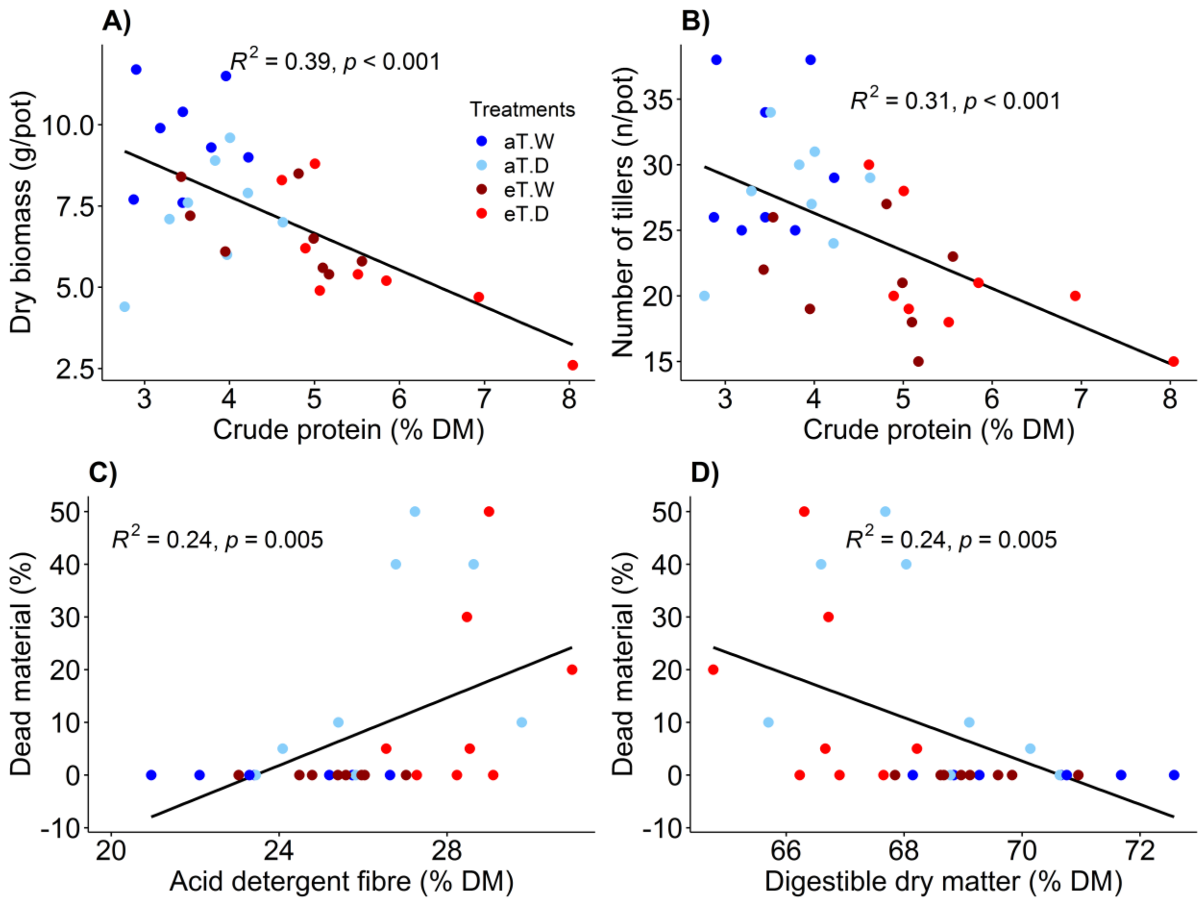


**Figure A1**. Correlations between A) plant dry biomass, morphological traits [B) number of tillers; C) and D) percentage of dead material] and nutritional composition [A) and B) crude protein; C) acid detergent fibre; D) digestible dry matter; all parameters as % of dry matter] for tall fescue (*Festuca arundinacea)* grown under different treatments (ambient, aT; elevated, eT, well-watered, W; droughted, D). Correlations were tested with Pearson correlation, with $R^2$ and *p* values shown in each panel.

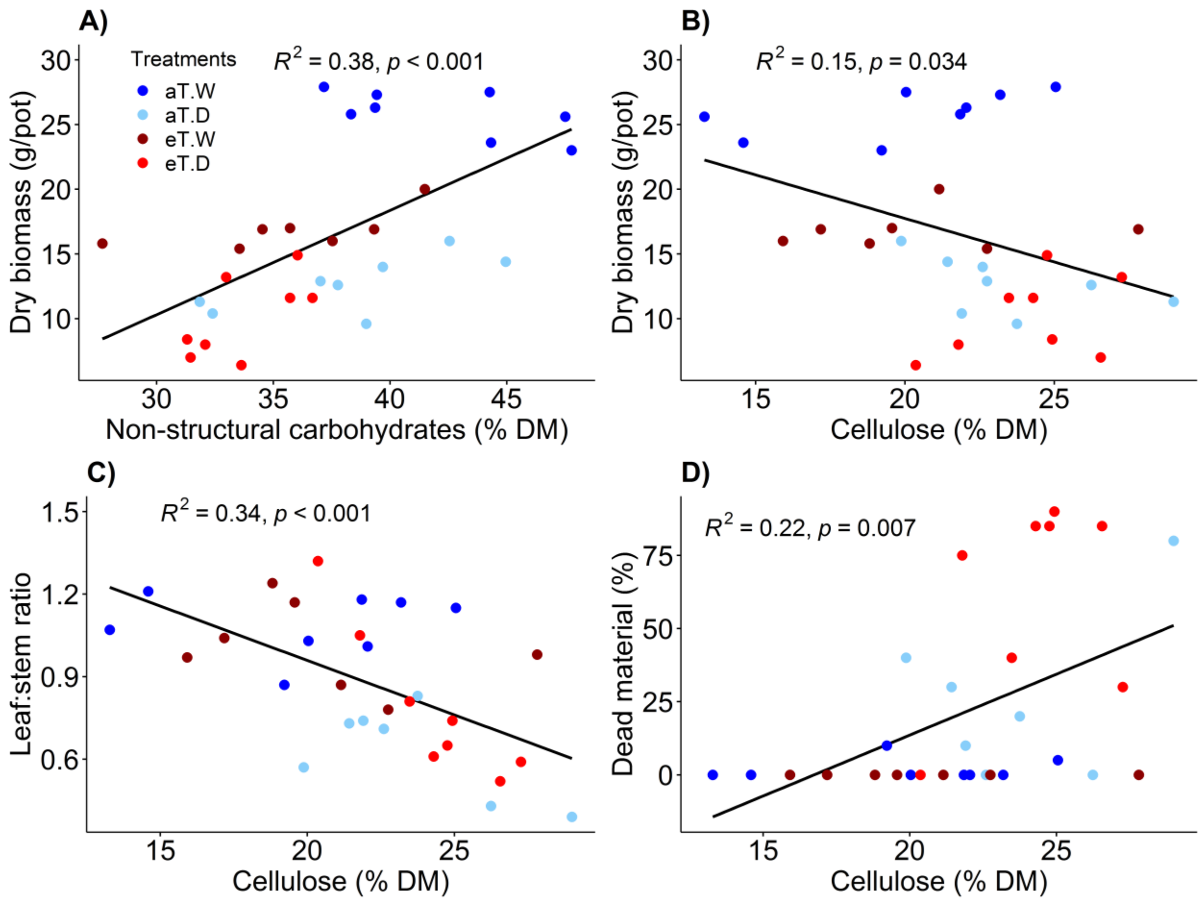


**Figure A2.** Correlations between plant dry biomass [A) and B)], morphological traits [C) leaf:stem ratio; D) percentage of dead material) and nutritional composition [A) non-structural carbohydrates; B), C) and D) cellulose; all parameters as % of dry matter) for lucerne (*Medicago sativa)* grown under different treatments (ambient, aT; elevated, eT, well-watered, W; droughted, D). Correlations were tested with Pearson correlation, with $R^2$ and $p$ values shown in each panel.
